# Modulating Glutamine Metabolism Reprograms Pro-Inflammatory Differentiation in Macrophages

**DOI:** 10.1101/2025.09.16.675216

**Authors:** Sumeng Qi, Jiawei Fan, Dao-Sian Wu, Sudipto Ganguly, Ie-Ming Shih, Tian-Li Wang

**Affiliations:** Graduate Program in Biochemistry and Molecular Biology, Bloomberg School of Public Health, Johns Hopkins University; Departments of Gynecology/Obstetrics, Oncology, and Pathology, School of Medicine, Johns Hopkins University

**Keywords:** Glutamine Metabolism, BMDM, Macrophage Differentiation, Pro-Inflammatory Genes, Tumor Microenvironment

## Abstract

Previous in vivo studies demonstrated that JHU083/DON, a glutamine analog drug, potently reprograms M1/M2 macrophages. To determine whether these effects are direct or indirect, we utilized an in vitro murine bone marrow–derived macrophage (BMDM) model, which recapitulates macrophage differentiation and polarization processes, to examine the impact of DON on the M1 macrophages. DON was applied during M1 differentiation or to fully polarized M1 macrophages, revealing that glutamine inhibition initially suppressed M1 activity but later enhanced it, resulting in sustained pro-inflammatory activation. Multi-omics analyses (bulk RNA-seq and LC-MS), time-course assays, and glutamine depletion experiments consistently suggested that prolonged glutamine inhibition elevates glutamine levels, which sustain pro-inflammatory gene transcription. In contrast, M2 and tumor-associated macrophages (TAM), which are immunosuppressive, were more susceptible to DON, leading to functional suppression. Collectively, our findings uncover stage-specific mechanisms by which glutamine inhibition modulates M1 polarization, offering a mechanistic rationale for therapeutic strategies that sustain pro-inflammatory, anti-tumor macrophage activity while concurrently suppressing immunosuppressive myeloid subsets in cancer.

## Introduction

Macrophages are critical components of the innate immune system, playing a central role in shaping the tumor microenvironment (TME) and adopting diverse phenotypes in response to environmental cues. Classical macrophage polarization is broadly categorized into pro-inflammatory M1 and anti-inflammatory M2 phenotypes. M1 macrophages exert anti-tumor effects by producing inflammatory cytokines (e.g., IL-1, IL-6, IL-12, TNF), enhancing antigen presentation (e.g., MHC-II), and stimulating cytotoxic immune responses.^1,2^ In contrast, M2 macrophages facilitate tumor progression by promoting angiogenesis and secreting anti-inflammatory cytokines (e.g., IL-10).^3^ In tumor tissues, the unique tumor microenvironment shapes the unique phenotype of tumor-associated macrophages (TAMs), which usually transform to the M2-like immunosuppressive phenotype and promote tumor progression and metastasis.^4–6^ Distinct metabolomic programs likely drive differentiation and functional polarization of M1 and M2 macrophages. However, due to the complexity of model systems and context-dependent phenotypes associated with various pharmacological drugs with distinct actions employed in the published research, the immunometabolism of anti-tumor macrophage activation and reprogramming remains to be explored.^6–8^

To address this, several methods can be employed. Single-cell RNA sequencing enables detailed analysis of gene expression at the individual cell level. Advanced imaging techniques, such as multiphoton microscopy, provide insights into spatial interactions between macrophages and tumor cells. Additionally, metabolic flux analysis can help identify the specific metabolic pathways involved in macrophage polarization within the tumor microenvironment. Key metabolites implicated include aKG, itaconate, glutamine, and succinate.

Glutamine, a non-essential amino acid, exerts distinct effects on M1 or M2 macrophage polarization by driving metabolic reprogramming.^9^ As a broadly active glutamine antagonist, 6-Diazo-5-oxo-L-norleucine (DON) inhibits multiple glutamine-utilizing enzymes. However, due to its toxicity to normal tissues, DON has limited clinical application.^10^ The development of DON prodrugs, JHU083 series, offers new hope for using glutamine metabolic inhibitors for cancer tissue-specific treatment. By chemically conjugating DON with chemical moieties such as leucine or the ethyl ester group, the DON prodrugs remain largely inert in circulatory systems and are predominantly activated by peptidase or esterase overexpressed in carcinoma tissues, leading to the release of active DON at the tumor site.^11^ This tumor-selective activation maximizes cytotoxicity within the tumor microenvironment while minimizing off-target effects on normal tissues.^11^ Both DON and JHU083 have been shown to play pivotal roles in suppressing tumor progression and remodeling the tumor immune microenvironment.^4,6,11–13^ In the endometrial and ovarian cancer model, DON/JHU083 has been reported to downregulate immunosuppressive immune cell populations.^4^ Recent studies in prostate and bladder cancer models, various in vivo cancer models have demonstrated that DON/JHU083, when used as a single agent, elicits potent antitumor efficacy.^6,13^ This effect may be partly attributed to pro-inflammatory macrophage reprogramming stimulated by DON/JHU083.^6,13^ For instance, by using ex vivo bone marrow macrophage cultures, we observed that glutamine inhibition not only suppressed M2-like macrophages but also enhanced M1-associated features, including increased MHC-II–mediated antigen presentation and IL-12 production.^4^

However, the precise mechanisms by which glutamine pharmacological inhibition reshapes the macrophage polarization program remain unclear. Given the robust and reproducible anti-tumor efficacy of DON/JHU083 and other glutamine inhibitors (e.g., CB-839) observed in animal models^14^, it is critical to gain molecular insights on how this novel class of drugs modulates immunometabolism to elicit anti-tumor immunity, resulting in superior cancer control over conventional glutamine metabolism inhibitors.

In the current study, we used an ex vivo macrophage differentiation model to systematically investigate stage-specific effects of glutamine inhibition via DON/JHU083 on M1 macrophage phenotypes. The study provides comprehensive transcriptomics and metabolomics pictures of DON/JHU083 in stimulating M1 macrophages at different developmental stages. The unexpected enhancement of glutamine and its metabolism by DON/JHU083, rather than inhibition in M1 macrophages, highlights the unique pharmacological profiles of this inhibitor class. This information is valuable for developing cancer therapeutic strategies that target unique metabolic programs in macrophages to enhance antitumor immunity.

## Materials and Methods

### Cell Culture and Macrophage Differentiation

Bone marrow cells were isolated from the tibia and femur of C57BL/6 mice and were cultured in Advanced DMEM/F12 Medium (Gibco) with 10% FBS, 1% Pen/Strep, 1% GlutaMax (Gibco), and m-CSF (50 ng/ml) in low-attachment culture plates for 5 days to generate bone marrow-derived macrophages (BMDM). For M1 macrophage polarization, two protocols were used: (1) Differentiation: cells were treated with Lipopolysaccharide (LPS, 10 ng/ml) for 4-24 hours, with or without the addition of 6-Diazo-5-oxo-L-norleucine (DON, 10 μM); (2) Fully-Polarized: cells were first stimulated with LPS (10 ng/ml) for 24 hours to achieve full M1 polarization, followed by treatment with or without DON (10 μM) for another 24 hours. For M2 macrophage differentiation, cells were treated with IL-4 (10 ng/ml) for 24 hours with or without DON (5 μM). Cells without M1 or M2 stimulation were designated as M0. The resulting macrophages were harvested for downstream analyses.

For glutamine depletion experiments, cells were cultured with or without 1% GlutaMax supplement (Gibco), representing glutamine-sufficient (GLN^+^) or glutamine-depleted (GLN^−^) conditions, respectively. For glucose modulation tests, glucose concentrations were adjusted to 25 mM (High Glucose) or 2.5 mM (Low Glucose) using Glucose solution (Gibco).

To generate tumor-associated macrophage (TAM)-like cells, ID8 ovarian cancer cells were cultured for 2 days, after which the culture supernatant was collected and combined with standard BMDM culture media in a 1:1 ratio. M0 BMDMs were subsequently incubated in this tumor-conditioned media for 24 hours. DON (10 μM) or vehicle control was added during this incubation phase.

RAW 264.7 cell line was cultured in Advanced DMEM medium (Gibco) with 10% FBS, 1% Pen/Strep, and 1% GlutaMax (Gibco). To induce M1 polarization, RAW 264.7 cells were treated with 10 ng/mL LPS for 16 hours. DON (10 μM) or vehicle control was added during the LPS incubation phase.

### qRT-PCR

Total RNA was extracted using the RNeasy Micro Kit (QIAGEN) following the manufacturer’s instructions. RNA concentration and purity were assessed using a NanoDrop spectrophotometer. Only samples with an OD260/280 and OD260/230 ratio ≥ 2.0 were used for downstream analyses. Purified RNA was stored at −80°C until further use.

Total RNA was reverse-transcribed into cDNA using a reverse transcription kit (Invitrogen) following the manufacturer’s instructions. Quantitative PCR was performed using SYBR Green Super Mix (BioRad) and gene-specific PCR primers (STAR Table). Gene expression levels were normalized to the expression of a housekeeping gene, Amyloid Protein Precursor (APP1), and cross-sample comparison was evaluated using the 2^−ΔΔCt method.

### Flow Cytometry

Approximately 1 × 106 cells were analyzed via flow cytometry for each sample. Cells were first incubated with Fixable Viability Dye (Invitrogen) for 15 minutes at room temperature, followed by Fc receptor blocking using the mouse Fc block reagent for 5 minutes at 4°C. Cells were then stained with fluorescent-labeled antibodies targeting surface markers, including CD45, CD68, CD301b, and MHC-II, for 30 minutes at 4°C in the dark. Samples were washed with 1X PBS 2 times. After washing, samples were analyzed on a CytoFLEX S flow cytometer (Beckman Coulter). Data were processed using CytExpert software (Beckman Coulter). Gating strategies are available upon request.

### Phagocytosis assay

M1-polarized BMDMs treated with DON or vehicle control were collected and seeded at a density of approximately 5 × 10^5^ cells per well in a 6-well plate with standard culture medium. Cells were then incubated with bioparticle beads (ThermoFisher) for 15 minutes at 37°C. Cells were subsequently harvested and analyzed by flow cytometry to determine the percentage of PE-positive, phagocytosis-active cells.

### Metabolomic Analysis by LC-MS

M1-polarized BMDMs treated with DON or vehicle control were collected for mass spectrometry analysis. Approximately 30 μL (∼10M cells) of cell suspension per sample was submitted to the Metabolomics Core Facility at Weill Cornell Medicine for liquid chromatography-mass–mass spectrometry (LC-MS) analysis.

### Invasion Assay

Transwell invasion assays were carried out utilizing 24-well Invasion Chambers with Matrigel Matrix (Fisher Scientific). TAMs or M1 BMDMs pretreated with DON or vehicle control were seeded into the lower chamber at a density of approximately 6 × 10^5^ cells per well. ID8 ovarian cancer cells (1 × 10^5^ cells) were seeded into the upper chamber (insert). Co-cultures were incubated for 24 hours. Afterwards, a cotton swab was used to remove non-invading cells, which remained on the surface of the upper chamber. Inserts were then fixed with methanol and stained with crystal violet. Photos were taken on the stained insert, and the relative cell numbers were quantified as the mean staining intensity using ImageJ.

### ELISA

Abcam’s ELISA kits were used to measure the concentrations of IL-1β and IL-6 in culture supernatants, following product recommendations. Cytokine concentrations were calculated based on standard curves generated using the kit-provided standards. OD450 was measured using a FLUOstar Omega microplate reader (BMG LABTECH; software version 6.20, firmware version 1.53).

### Luminescence-Based Metabolite Assay

Cellular metabolites, including pyruvate, lactate, glutathione (GSH), and NAD, were quantified using luminescence-based assay kits from Promega. Cell pellets were collected at the end of the experiment and processed following the manufacturer’s protocols. Metabolite concentrations were calculated based on standard curves generated using the kit-provided standards. Luminescence was measured using a FLUOstar Omega microplate reader (BMG LABTECH; software version 6.20, firmware version 1.53).

### Cell Viability Assay

Cell viability was assessed using the CellTiter-Blue® Cell Viability Assay (Promega). M0 BMDMs were treated with LPS (10 ng/mL) for 16 hours or IL-4 (10 ng/mL) for 24 hours to polarize them into M1 or M2 macrophages. M1 or M2 macrophages were seeded into 96-well plates and treated with a gradient of DON (0–40 μM) for 24 hours. After treatment, the medium was removed and replaced with fresh medium containing CellTiter-Blue reagent at a 1:6 ratio. Cells were incubated for 3 hours at 37 °C. Fluorescence intensity was measured using a FLUOstar Omega microplate reader (BMG LABTECH; software version 6.20, firmware version 1.53). Fluorescence readings were normalized to the 0 μM DON condition within each group, which was set as 100% viability for the respective cell type.

Apoptosis was assessed using the Caspase-Glo® 3/7 Assay (Promega) according to the manufacturer’s protocol. M0 BMDMs were polarized into M1 or M2 macrophages as described above, seeded into 96-well plates, and treated with a gradient of DON (0–20 μM) for 24 hours. Luminescence was measured using a FLUOstar Omega microplate reader, and caspase 3/7 activity values were normalized to the 0 μM DON condition within each group, which was set as 1.0.

### RNA sequencing and Data analysis

Sequencing was performed by Novogene Corporation (USA) on an Illumina NovaSeq X Plus platform using paired-end 150 bp (PE150) sequencing, resulting in approximately 6 Gb of raw data per sample. Eight samples were sequenced, with two replicates for each group: Ctrl_4h, DON_4h, Ctrl_16h, DON_16h. Raw reads were quality-checked and trimmed to remove adaptor contamination, low-quality bases (Q < 5), and reads containing >10% ambiguous bases (N). Clean reads were aligned to the mouse reference genome (GRCm39/mm39) with HISAT2. Gene-level expression was quantified using featureCounts or HTSeq.

Differential gene expression analysis was conducted using the DESeq2 package in R. Genes with an adjusted p-value < 0.05 and log_2_ change ≥ 1 were considered significantly differentially expressed. Genes with low expression (mean normalized counts < 10) across all samples were filtered out before the analysis. R was used to perform RNA-seq downstream data analyses, including differential expression visualization (volcano plots and heatmaps), KEGG pathway enrichment analysis, gene set enrichment analysis (GSEA), and expression overlap analysis using Venn diagrams. ***Ingenuity Pathway Analysis (IPA)*** was used to interpret differentially expressed genes (DEGs) identified by DESeq2. For each comparison (Treated vs. Control at 4h and 16h), filtered DEGs (adjusted p-value < 0.05 and |log_2_ fold change| ≥ 1) were analyzed by IPA’s Core Analysis. Genes with low expression (mean normalized counts < 10) across all samples were removed before analysis. Canonical pathways, upstream regulators, and regulatory interaction networks were assessed.

### Statistical analysis

All plots and statistical analyses were generated using GraphPad Prism software. To compare the DON treatment with the control group, unpaired Student’s t-tests were utilized. All statistical tests were performed as two-tailed, and p-values of less than 0.05 were considered significant. Each condition was tested in three independent biological replicates, and two to three technical replicates were included per biological replicate to minimize experimental variability.

## Results

### Glutamine antagonism differentially affects tumor-suppressive functions and pro-inflammatory phenotype of M1 macrophages during polarization

To investigate the impact of glutamine metabolism on M1 macrophage activation, we established an ex vivo primary culture model that allows for stage-specific analysis of M1 differentiation and polarization. As illustrated in **Figure 1A**, mouse bone marrow cells were differentiated into bone marrow-derived macrophages (BMDMs) using M-CSF. These cells were designated as M0 macrophages, which lack the surface markers characteristic of M1 and M2 macrophages. M1 polarization was induced by LPS (10 ng/mL) for 24 hours to achieve full polarization.

**Figure 1.**
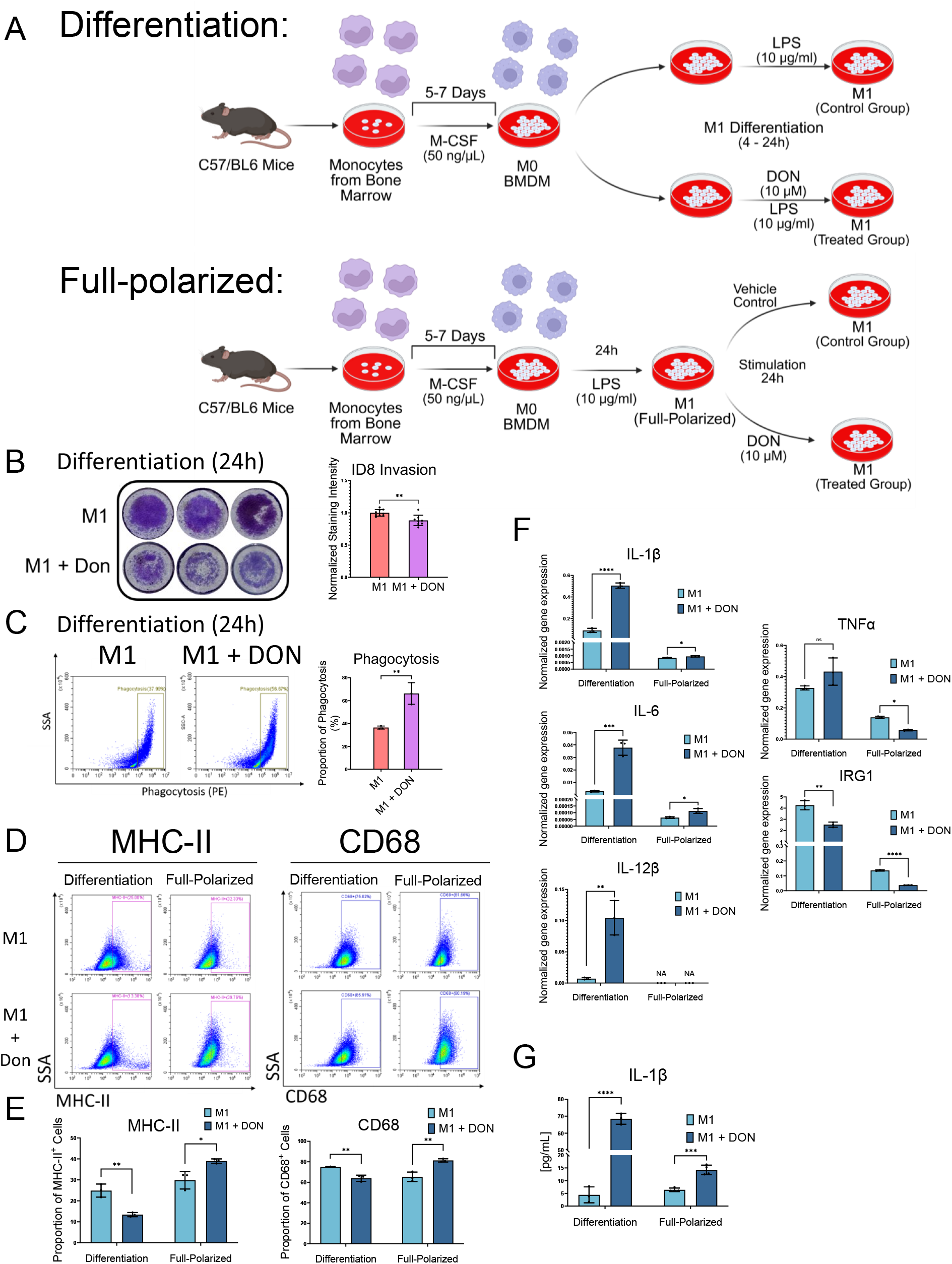
The glutamine metabolism drug DON stimulates M1 macrophage function and phenotype. (A) Experimental scheme for M1 macrophage polarization with or without DON treatment. Bone marrow cells were harvested from the tibia and femur of C57BL/6 mice and stimulated with M-CSF (50 ng/mL) for 5–7 days to produce bone marrow-derived macrophages (BMDMs). To assess DON’s effect on M1 differentiation stage, BMDMs were treated with LPS (10 ng/mL) for 4-24 hours, with or without DON (10 μM) added simultaneously. To assess DON’s effect on M1 polarization, BMDMs were first stimulated with LPS (10 ng/mL) for 24 hours to achieve full M1 polarization, followed by treatment with or without DON (10 μM) for an additional 24 hours. Resulting macrophages were used for downstream analysis. (B) Transwell invasion assay of ID8 ovarian cancer cells co-cultured with M1-differentiated BMDMs. M1 macrophages following the treatment protocol described in (A) were co-cultured with ID8 cancer cells for 24 hours (in the absence of DON) for a transwell invasion assay. Invaded cells were stained with crystal violet; the relative cell number was determined by the mean staining intensity. (C) Phagocytosis assay using pH-sensitive PE-labeled E. coli bioparticles. M1-differentiated BMDMs following treatment protocol described in (A) were incubated with PE-labeled E. coli bioparticles for 15 minutes. Phagocytic activity was assessed by flow cytometry and quantified based on the proportion of PE^+^ cells versus total cells. (D) Flow cytometry analysis of MHC-II and CD68 expression in BMDMs under both differentiation and full-polarized conditions, with or without DON treatment. (E) Quantification of the flow cytometry results shown in (D), representing the percentage of MHC-II^+^ and CD68^+^ cells in each group. (F) qPCR analysis of M1-associated genes (IL-1β, IL-6, TNFα, IL-12β, and IRG1) in BMDMs treated with or without DON under differentiation and full-polarized conditions. (G) ELISA measurement of IL-1β protein levels in the culture supernatant of M1-polarized BMDMs treated with or without DON under both differentiation and full-polarized conditions. Data represent mean ± SEM from at least three independent experiments. Statistical significance was assessed by unpaired two-tailed Student’s t-test (p < 0.05; p < 0.01; p < 0.001; p < 0.0001; ns, not significant).

In the M1 differentiation model, DON, a broadly active glutamine antagonist, was applied concurrently with LPS to examine its effects on the M0 to M1 transition (**Figure 1A, top**). In the M1 fully polarized model, BMDMs were first stimulated with LPS to achieve full polarization, followed by the addition of DON to target already polarized M1 macrophages (**Figure 1A, bottom**).

We then evaluated the ability of DON-modulated M1 macrophages to promote tumor invasion. M0 BMDMs were treated with LPS following the differentiation protocol in Figure 1A, then seeded into 24-well plates. ID8 ovarian cancer cells were seeded in the upper Matrigel-coated chambers and co-cultured with the BMDMs. The results demonstrated that the invasion of ID8 ovarian cancer cells was significantly suppressed when co-cultured with DON-treated M1 macrophages. A phagocytosis assay was also conducted using the PE-conjugated E. coli bioparticles and quantified for PE-positive cells by flow cytometry. In the M1 differentiation model, a higher proportion of PE-positive cells was detected in the DON-treated macrophages, indicating enhanced phagocytic activity (**Figure 1C**). In contrast, in the M1 full-polarized model, DON treatment did not significantly affect the invasion-suppressive or phagocytic capacities of M1 macrophages (**Supplementary Figure 1A &1B**).

We next investigated the expression of M1-specific markers in both differentiation and fully-polarization models to assess the impact of DON treatment at these two stages. Flow cytometry analysis of MHC-II^+^ and CD68^+^ surface marker expression revealed that DON had opposite effects depending on the treatment stage **(Figure 1D– E)**. In the differentiation model, DON decreased the proportion of MHC-II^+^ and CD68^+^ macrophages compared to the control. In the fully polarized model, DON increased the expression of both MHC-II and CD68 (**Figure 1D–E)**.

Expression of the M1-associated cytokines (IL-1β, IL-6, IL-12β), classical M1 marker (TNFα), and IRG1 were evaluated by qRT-PCR. Notably, IRG1 was included based on its reported role in suppressing the M1 inflammatory response through its enzymatic product, itaconate.^7^ In the differentiation model, DON treatment markedly increased the expression of IL-1β, IL-6, and IL-12β while reducing IRG1 expression, indicating a shift toward a pro-inflammatory phenotype (**Figure 1F**). TNFα showed no significant increase, possibly reflecting gene-specific responses to glutamine metabolism. In the fully polarized model, baseline levels of the M1-associated cytokines are significantly lower than in the differentiation stage, suggesting limited pro-inflammatory transcriptional activity at this stage. Modest increases in IL-1β and IL-6 were detected in the DON-treated group compared to the vehicle-treated group (**Figure 1F**). IL-1β, a hallmark cytokine of M1 macrophages involved in pro-inflammatory responses, was selected for ELISA analysis. Results showed that IL-1β protein was significantly elevated by DON in the differentiation model **(Figure 1G)**, supporting stage-specific cytokine production.

To confirm the generality of the findings, we examined the expression of M1-associated cytokines and surface markers in RAW264.7 cells, a murine macrophage cell line. The results closely mirrored those observed in primary BMDMs, reinforcing the robustness of the stage-dependent effects of glutamine antagonism **(Supplementary Figure 2A-C)**.

### DON-Mediated Transcriptomic Reprogramming Promotes M1 Macrophage Activation

We next performed bulk RNA sequencing to identify potential mechanisms underlying glutamine-mediated M1 macrophage reprogramming. BMDMs were treated with DON for 4 hr and 16 hr following the protocol described in Figure 1A. DMSO (vehicle control) was applied to BMDMs for the same duration. The 4 hr treatment group and the 16 hr treatment group exhibited differential gene expression patterns (**Figure 2A**). When comparing the DON-treated and DMSO control-treated groups, there are 1293 differentially expressed genes (DEG) (|LogFC| > 1 & p-value < 0.05) at 4 hr incubation and 1380 DEG at 16 hr incubation time (**Figure 2B**), among which 277 genes overlap between the two time points (**Figure 2C**). **Figure 2D** shows the overlap of highly expressed genes (log_2_ [counts/gene number] > 1) among the control (4 h, 16 h) and DON-treated (4 h, 16 h) groups. Apoptosis-related genes (e.g., Bcl2l11) and inflammation-related genes (e.g., Rgs1, Ccrl2, and Tnfsf9) were upregulated in the 4 hr treatment group. In contrast, M1-associated cytokines (e.g., IL-12b) and guanylate-binding protein (GBP) family genes were upregulated at the 16 hr time point (**Figure 2E**). M2-associated markers such as Ccl2 and Ccl5 are downregulated at 16 hr DON treatment condition (**Figure 2B**).

**Figure 2.**
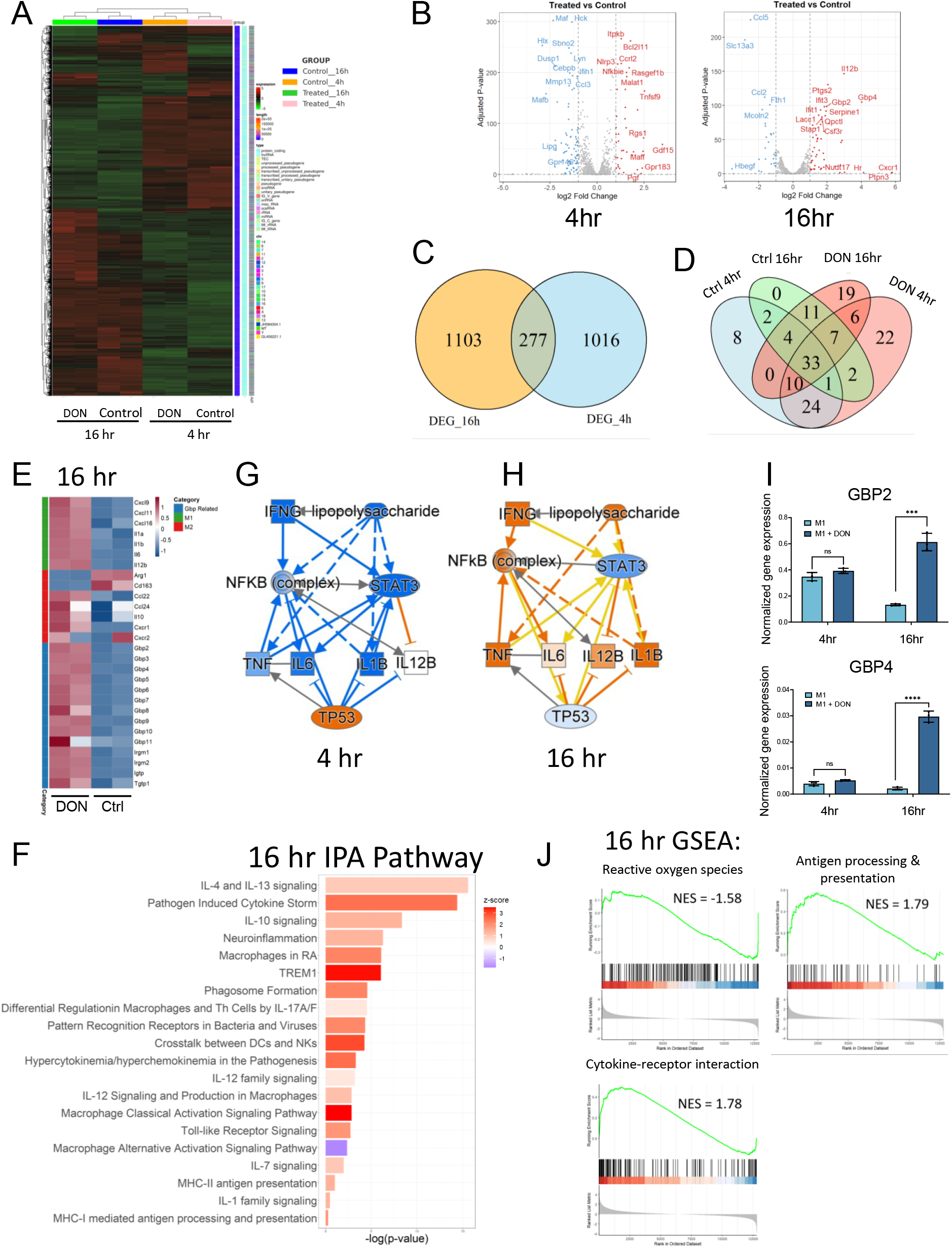
DON-Mediated Transcriptomic Reprogramming Drives M1 Macrophage Activation. (A) Heatmap showing differentially expressed genes (DEGs) identified by bulk RNA-seq in M1-differentiated cells treated with DON or vehicle during 4 hr and 16 hr LPS stimulation. (B) Volcano plots showing differentially expressed genes between DON and vehicle control-treated group at each time point. (C) Venn diagram showing the overlap of differentially expressed genes at 4 hrs and 16 hrs of LPS-stimulated differentiation stage. DEGs were defined as genes with |log_2_ fold change| > 1 and adjusted *p*-value < 0.05. (D) Venn diagram of highly expressed genes across the four groups. Genes with log_2_(counts/gene number) > 1 were considered highly expressed. (E) Heatmaps of representative M1/M2 markers and GBP family genes. (F) Ingenuity Pathway Analysis (IPA) of DEGs at 16 hours, highlighting immune- and macrophage-related pathways. (G, H) IPA upstream regulator analysis at 4 h (G) and 16 h (H). Node color indicates predicted activation state (orange: activated; blue: inhibited). (I) qPCR validation of selected GBP genes at 4 h and 16 h. (^*^ p<0.05; ^**^ p<0.01; ^***^ p<0.001; ^****^ p<0.0001; ns, not significant). (J) Gene Set Enrichment Analysis (GSEA) based on bulk RNA-seq data comparing treated versus control macrophages at 16 hours. Normalized Enrichment Score (NES) and adjusted p-values (P.adjust) are as follows: Chemical carcinogenesis – reactive oxygen species: NES = –1.58, P.adjust = 4.62e-03; Cytokine– cytokine receptor interaction: NES = 1.78, P.adjust = 8.58e-04; Antigen processing and presentation: NES = 1.79, P.adjust = 5.18e-03

KEGG pathway analysis showed immune-related signaling pathways, including the NF-κB and the JAK-STAT, were enriched among the differentially expressed genes, suggesting a changed immune response and functions of the macrophage (**Supplementary Figure 4**). At the 16 hr treatment condition, we observed an overall M1-activated cytokine expression pattern. Ingenuity Pathway Analysis (IPA) of the DEG genes **(Table S1)** revealed up-regulation of the macrophage classical activation signaling pathway, pathogen-induced cytokine storm, trem1 pathway, and IL-12-related pathway, suggesting the enhanced pro-inflammatory macrophage activity (**Figure 2F, Table S2**). In addition, LPS-mediated signaling, TNF signaling, and IFN-gamma signaling, which were down-regulated in the 4 hr DON treatment group, were up-regulated in the 16 hr DON treatment group (**Figures 2G-H**).

qRT-PCR was performed and confirmed the expression pattern of Gbp2 and Gbp4 at different time points (**Figure 2I**). We also used Gene Set Enrichment Analysis (GSEA) to compare differentially expressed genes between DON and DMSO control treatments. We discovered that in the 16 hr treatment group, gene sets related to cytokine-receptor interaction and antigen processing and presentation were significantly upregulated in the DON-treated group, while the reactive oxygen species gene set was significantly downregulated (**Figure 2J**). On the other hand, the 4 hr treatment group showed upregulation of stress-associated pathways such as p53 and apoptosis **(Supplementary Figure 5)**. The findings collectively indicated that long-term (16 hr) DON treatment enhances M1 macrophage activation.

### Metabolic Reprogramming in M1 Macrophages Induced by DON Treatment

To assess metabolites induced by DON treatment that may underlie the pro-inflammatory M1 phenotypes, we performed liquid chromatography-mass spectrometry (LC-MS) on BMDMs after 16 hr of DON or vehicle control treatment. Metabolites showing differential abundances between DON and vehicle control treatment are summarized in **Figure 3A** and **Table S3**. We examined glycolysis, TCA cycle, and glutamine metabolism pathways as potential metabolic regulators of macrophage function. The glucose level was upregulated, while glycolysis upstream metabolites, G6P and F1,6BP, were downregulated (**Figure 3B-C**). Lactate, the end product of glycolysis and important for M2 macrophage polarization, was downregulated. Most TCA cycle components showed no significant changes (**Figure 3D**). Meanwhile, L-glutamine, L-glutamate, and glutamine downstream metabolites, including NAD, NADH, NADP, NADPH, and GSH, were significantly upregulated (**Figure 3D, 3H**). Luminescence-based metabolite assays were performed and confirmed the downregulation of lactate as well as the upregulation of GSH and NAD by DON treatment (**Figure 3E**). Additionally, L-Aspartic acid and L-Arginine were significantly upregulated, whereas Ornithine was reduced (**Figure 3F-G**). In nucleotide metabolism, pyrimidines were elevated, and inosinic acid, the first nucleotide formed during the synthesis of purine nucleotides, was upregulated (**Figure 3I, Table S3**). Increased ratio of glutathione (GSH) and GSH/oxidized glutathione (GSSG), suggesting a more reductive environment for macrophages after DON treatment. To assess whether gene expression of enzymes in metabolic pathways confers the observed metabolite alterations, we performed gene set enrichment analysis (GSEA) and found that glutathione metabolism gene set was upregulated, whereas glycolysis and oxidative phosphorylation gene sets were repressed (**Figure 3J**). These findings suggest that DON treatment induces metabolic changes that support a pro-inflammatory M1 macrophage phenotype.

**Figure 3.**
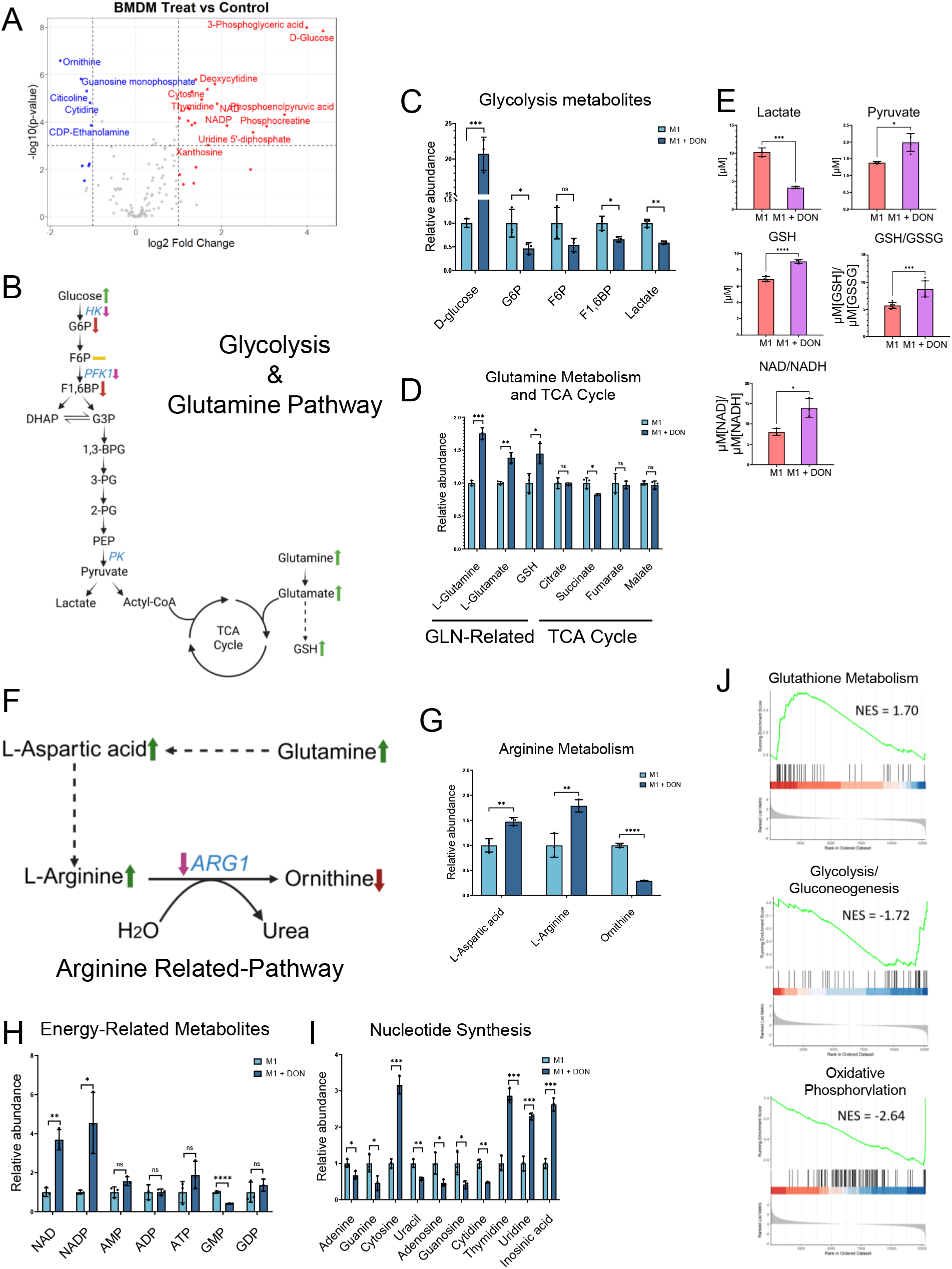
Metabolic Alterations in M1 Macrophages Induced by DON Treatment. (A) A volcano plot depicting the difference in metabolite abundance between DON-treated and control-treated M1-macrophages, differentiated after 16 hours. Global metabolites were analyzed by the LC-MS. (B) Schematic of glycolysis, the TCA cycle, and glutamine metabolism. Key metabolites (from LC-MS and luminescence-based assays) and relevant enzymes (from RNA-seq) are annotated with expression changes between DON-treated vs. control group. (C) Relative abundance of glycolytic metabolites measured by LC-MS. (D) Relative abundance of glutamine- and TCA cycle– related metabolites measured by LC-MS. (E) Quantification of lactate, pyruvate, GSH, GSH/GSSG, and NAD/NADH levels using luminescence-based enzymatic assays (ELISA). (F) Schematic of arginine-related pathway, with metabolite changes derived from LC-MS and gene expression changes of key enzymes obtained from RNA-seq. (G) Relative abundance of arginine-related metabolites measured by LC-MS. (H) Relative levels of energy-related metabolites (e.g., NAD, NADP, ATP, AMP, etc.) measured by LC-MS. (I) Relative abundance of nucleotide synthesis intermediates measured by LC-MS. (J) Gene Set Enrichment Analysis (GSEA) based on bulk RNA-seq data comparing treated versus control macrophages at 16 hours. Normalized Enrichment Score (NES) and adjusted p-values (P.adjust) are as follows: Glutathione metabolism: NES = 1.70, P.adjust = 3.59e-02; Glycolysis/Gluconeogenesis: NES = –1.72, P.adjust = 3.37e-02; Oxidative phosphorylation: NES = –2.64, P.adjust = 3.31e-08 Data represent mean ± SEM. p < 0.05; p < 0.01; p < 0.001; p < 0.0001 by unpaired two-tailed Student’s t-test.

### DON Treatment Stimulates Glutamine-Dependent, Pro-Inflammatory Macrophage Phenotypes

To evaluate temporal expression changes in detail, we performed a time-course expression analysis of M1-related genes during differentiation at 4, 8, 12, and 16 hours using qRT-PCR **(Figure 4A)**. In the control M1 differentiation condition, most pro-inflammatory cytokines exhibited a sharp and transient upregulation, peaking within the first 4–8 hours of LPS stimulation, followed by a rapid decline by 16 hours **(Figure 4A)**. This pattern suggests that the expression of M1-associated genes is tightly regulated and temporally restricted during M1-differentiation stages.

**Figure 4.**
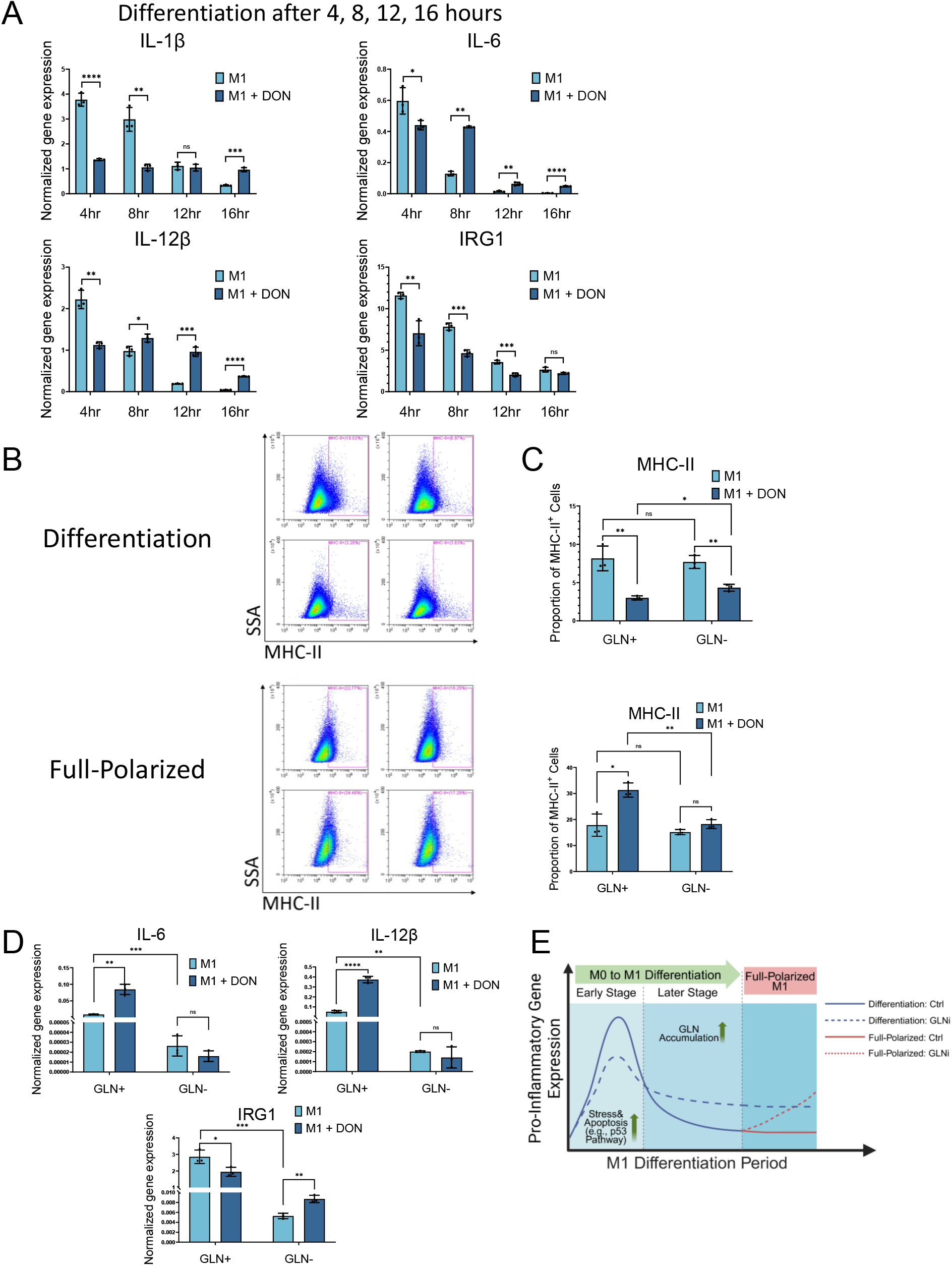
DON Treatment Stimulates Glutamine-Dependent, Pro-Inflammatory M1 Macrophage Phenotypes. (A) Time-course analysis of M1-associated gene expression during the differentiation stage by qPCR. BMDMs were treated with LPS alone or LPS plus DON for 4, 8, 12, or 16 hours. (B) Flow cytometry analysis of MHC-II expression under glutamine-repleted (GLN^+^) and glutamine-depleted (GLN^−^) conditions during both the differentiation and full-polarized stages. For the differentiation condition, BMDMs were stimulated with LPS for 16 hours in either GLN^+^ or GLN^−^ medium, with or without DON (10 μM). For the full-polarized condition, BMDMs were first stimulated with LPS for 24 hours in GLN^+^ medium, then cultured for an additional 16 hours in either GLN^+^ or GLN^−^ medium, with or without DON (10 μM). Comparisons were made between control and DON-treated groups within each condition. Up: differentiation model; down: full-polarized model. (C) Quantification of MHC-II^+^ cell proportions shown in (B). Up: differentiation model; down: full-polarized model. (D) qPCR analysis of M1-associated gene expression (IL-6, IL-12β, and IRG1) under glutamine-replete (GLN^+^) and glutamine-depleted (GLN^−^) conditions in the M1 differentiation model (16 hr). (E) Schematic model illustrating the stage-dependent regulation of pro-inflammatory gene expression by glutamine inhibition. The model depicts changes in pro-inflammatory gene expression during M0 to M1 differentiation (both early and late stages) and in fully polarized M1 macrophages under control and glutamine inhibition (GLNi) conditions. Data represent mean ± SEM from at least three independent experiments. Statistical significance was assessed by unpaired two-tailed Student’s t-test (p < 0.05; p < 0.01; p < 0.001; p < 0.0001; ns, not significant).

In the DON-treated M1 differentiation condition, expression of IL-1β, IL-6, and IL-12β was suppressed at the early time point (4 hr), but sustained at relatively higher levels during the mid-to-late differentiation stages (8–16 h) compared to the control **(Figure 4A)**. For IRG1, DON treatment consistently suppressed its expression across all time points, indicating a pro-inflammatory shift condition.

To test if extracellular glutamine is a critical source for promoting M1 macrophage differentiation, we cultured BMDMs in the glutamine-depleted (GLN^−^) or glutamine-replete (GLN^+^) medium and examined M1-associated MHC-II in both the differentiation and full-polarized conditions. As shown in **Figure 4B–C**, in the full-polarized model, DON significantly increased MHC-II expression in glutamine-replete (GLN^+^) medium, but this effect was nearly abolished in the glutamine-depleted (GLN^−^) conditions. In contrast, in the differentiation stage, DON consistently reduced MHC-II expression regardless of glutamine availability.

We further tested the extracellular glutamine dependency of M1 cytokines stimulated by DON at the 16-hour differentiation stage **(Figure 4D)**. Under GLN^−^ conditions, DON failed to upregulate IL-6 and IL-12β expression, indicating that the upregulation of pro-inflammatory cytokines requires extracellular glutamine. Additionally, the DON-induced suppression of IRG1 observed in GLN^+^ condition was eliminated in the GLN^−^ condition. In the short-term (4 h) differentiation model, glutamine-depleted conditions (GLN^−^) largely failed to abolish the DON-induced suppression of M1 cytokine transcription **(Supplementary Figure 6)**. These findings suggest that glutamine dependency differs between early and late stages of M1 differentiation, with only late-stage cells relying on an exogenous glutamine source for pro-inflammatory cytokine production.

Lastly, although according to LC-MS assays, glucose levels were elevated in DON-treated BMDMs **(Figure 3A–C)**, glucose depletion experiments revealed that DON’s effects on M1 polarization were not affected by glucose availability **(Supplementary Figure 7)**. This suggests that glutamine—but not glucose—plays a critical regulatory role in this process.

### M2 macrophages are more vulnerable to glutamine blockade

Next, to investigate the role of glutamine metabolism in M2 macrophage polarization, a BMDM-derived M2 macrophage model was employed. Bone marrow cells from C57/BL6 mice were cultured with M-CSF to induce M0 macrophage differentiation, followed by IL-4 treatment for 24 hours to promote M2 differentiation **(Figure 5A)**. DON was introduced during the IL-4 stimulation phase to assess its effects on M2 differentiation. qRT-PCR analysis showed a significant reduction in the expression of M2-associated genes, including Arg1, CCL22, IL-10, and Retnla, in the DON-treated group (**Figure 5B**). Flow cytometry further revealed a decrease in the CD301b-positive M2 cell population (**Figures 5C & 5D**). suggesting that DON disrupts the M0 to M2 differentiation process and suppresses the expression of M2 immunosuppressive cytokines.

**Figure 5.**
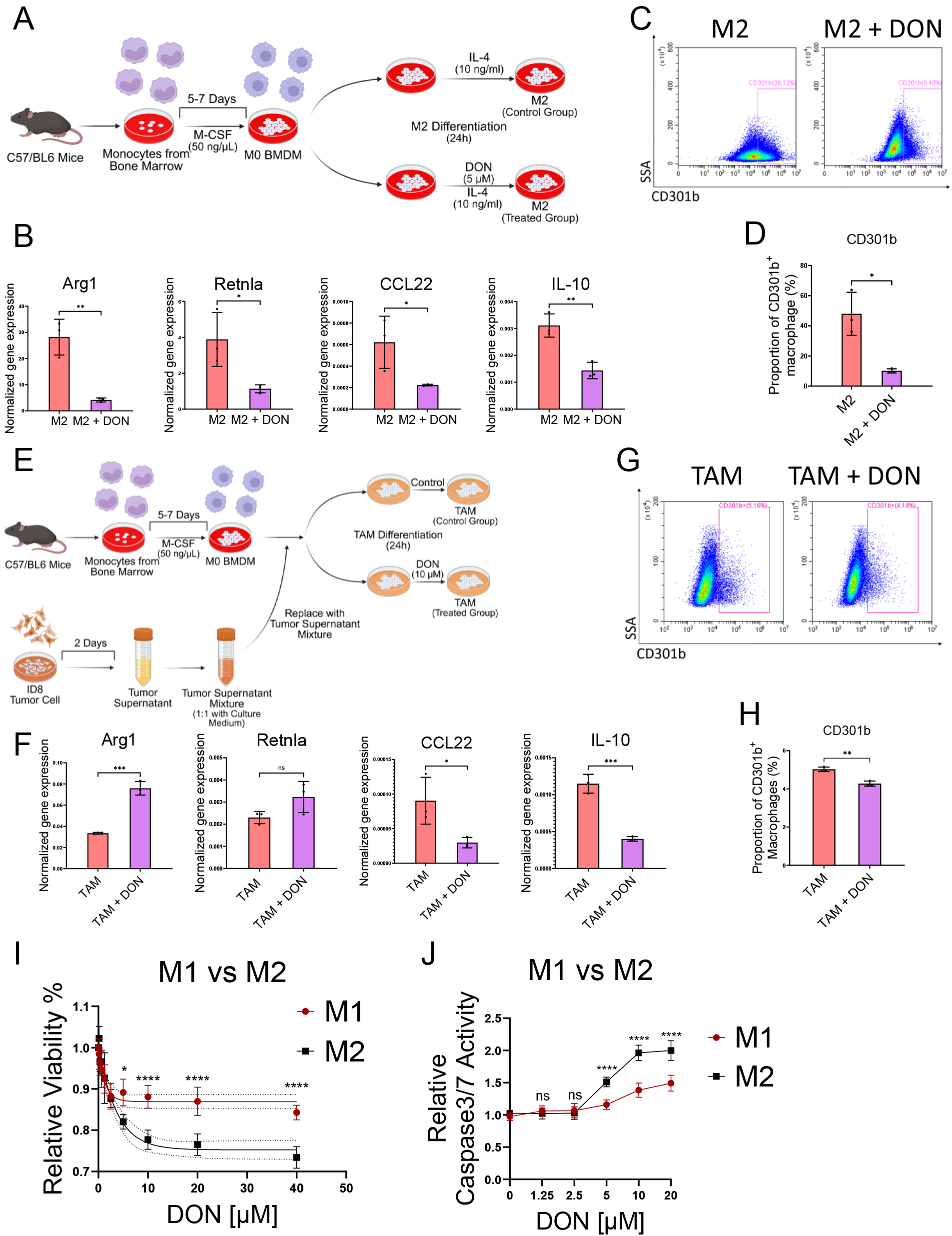
M2 macrophages are vulnerable to treatment with the glutamine metabolism drug, DON. (A) Experimental scheme for M2 macrophage polarization. BMDMs were treated with IL-4 (10 ng/mL) for 24 hours, with or without DON (5 μM), to induce M2 differentiation. (B) Relative expression of M2-associated genes (Arg1, CCL22, IL-10, Retnla) was assessed by qPCR after DON or DMSO treatment. (C) Representative flow cytometry plots showing CD301b expression in M2 macrophages prepared by the protocol described in (A). (D) Quantification of CD301b^+^ BMDM populations from (C). (E) Experimental scheme for producing TAM-like macrophages. BMDMs were incubated with tumor cell– conditioned medium (50%) for 24 hours, with or without DON. (F) qRT-PCR analysis of TAM-associated gene expression (IL-10, Arg1, CCL22, Retnla). (G) Flow cytometry plots showing CD301b expression in TAM-like macrophages. (H) Quantification of CD301b^+^ BMDM populations from (G). (I) Viability assay comparing the sensitivity of M1 and M2 macrophages to DON. Fully polarized M1 and M2 macrophages were seeded in 96-well plates and treated with increasing concentrations of DON (0–40 μM) for 24 hours. Cell viability was measured using the CellTiter-Blue assay and normalized to the 0 μM DON condition (set as 100%). (J) Apoptosis assay comparing the sensitivity of M1 and M2 macrophages to DON. Fully polarized M1 and M2 macrophages were seeded in 96-well plates and treated with increasing concentrations of DON (0–20 μM) for 24 hours. Apoptosis was measured using the Caspase-Glo® 3/7 assay and normalized to the 0 μM DON condition (set as 1.0). Data represent mean ± SEM from at least three independent experiments. Statistical significance was assessed using unpaired two-tailed Student’s t-test for panels A–H, and two-way ANOVA with Šidák’s multiple comparisons test for panel I–J (^*^p < 0.05; ^**^p < 0.01; ^***^p < 0.001; ^*^p < 0.0001; ns, not significant).

We further assessed the effect of DON in a tumor-associated macrophage (TAM) model (**Figure 5E**). In this model, M0 BMDMs were exposed to tumor cell-conditioned medium (supernatant from ID8 tumor cells cultured for 48 hours) following the protocol outlined in **Figure 5E**. We observed that tumor cell-conditioned medium promoted M0 BMDMs differentiated into an M2-like TAM phenotype (**Figure 5F**). DON treatment during TAM polarization resulted in a mixed response, while M2 markers Arg1 and Retnla were enhanced, functional cytokines CCL22 and IL-10 were suppressed (**Figure 5F**). Flow cytometry confirmed a reduction in the M2 CD301b+ population in the DON-treated condition (**Figure 5G-H**), indicating that DON impairs M2 function in TAMs, potentially disrupting the immunosuppressive tumor microenvironment.

To explore the differential effects of DON on M1 and M2 macrophages, cell viability and apoptosis assays were conducted on fully polarized M1 and M2 macrophages treated with a gradient of DON (0–40 μM). M2 macrophages exhibited greater sensitivity to DON, as indicated by a greater reduction in viability (**Figure 5I)**. Similarly, apoptosis, measured by Caspase-Glo® 3/7 activity, was significantly higher in M2 macrophages compared to M1 macrophages **(Figure 5J)**. These results suggest that M2 macrophages are more susceptible to glutamine-targeting agents, resulting in sustained suppression of their functional phenotype.

## Discussion

Previous studies from our group and others have demonstrated that JHU083/DON treatment elicited potent anti-tumor immunity in murine tumor models. Notably, JHU083-treated tumor models exhibited significant macrophage reprogramming, enhancing the M1/M2 ratios.^4^ However, the detailed mechanisms underlying this reprogramming remain unclear. This study aims to determine whether DON/JHU083 treatment directly influences macrophage transcriptional programs and stimulates M1 functions. To address this, we employed an ex vivo culture model of M1 macrophage differentiation and polarization to evaluate the direct pharmacological effects of DON. We found that DON incubation (16 hr) directly promoted M1-associated pro-inflammatory functions, including the inhibition of tumor invasion and stimulation of phagocytosis. Additionally, genome-wide RNA-seq analysis revealed distinct temporal transcriptomic alterations.

Concurrently performed metabolomic profiling showed that DON treatment significantly increased intracellular glutamine and its downstream metabolites, including glutamate, GSH, L-aspartate/L-arginine, and nucleotides in M1 macrophages. Intracellular glucose was also elevated. The unexpected findings align with previous metabolic studies on T cells, where increased levels of glutamine, glutamate, and glucose were reported following JHU083/DON treatment.^6,11^ This metabolic pattern suggests that the rise in intracellular glutamine after DON treatment is unlikely to result solely from the enzymatic blockage of glutaminase (GLS). Instead, an elevated influx of metabolites, such as those facilitated by phagocytosis or membrane transporters, may contribute to the observed findings. Supporting this interpretation, we observed enhanced phagocytic activity in M1 macrophages following DON treatment.

The elevation of glutamine metabolism elicited by DON in M1 macrophages was further supported by increases in energy-related metabolites, including ATP, NAD, and NADP. The findings suggest that enhanced glutamine utilization by DON can sustain cellular bioenergetics, maintaining the energy supply necessary for pro-inflammatory functions.

Amino acids, such as L-aspartic acid and L-arginine, were also elevated in the DON-treated BMDMs, which could similarly be attributed to increased glutamine utilization. On the other hand, the expression of arginase 1 (ARG1), which catalyzes the conversion of L-arginine to ornithine, was suppressed by DON. Because ARG1 is a representative M2 marker, its suppression could translate to a reduced immunosuppressive phenotype observed in the in vivo studies applying JHU083/DON.^4,6^

In contrast to elevated glutamine metabolism following DON treatment in M1 macrophages, glycolysis was suppressed, as evidenced by a marked reduction in glycolysis intermediates and end products, such as G6P, F1,6 BP, and lactate. This was accompanied by a decrease in the expression of the rate-limiting enzymes, HK and PFK-1, of the glycolysis pathway. The results suggest that glycolysis is not essential for the differentiation or activation of M1 macrophages, which differs from M2-like TAMs, where glycolysis is often required to achieve full M2 function.^5,9^

Our study offers new insights into how broad glutamine inhibition affects the phenotype and function of M1 macrophages. Nevertheless, certain limitations should be acknowledged. First, we used an ex vivo BMDM model to focus on mechanistic investigations. However, in the in vivo contexts, macrophages are likely subject to a multitude of factors, such as nutrient competition within the TME, which influence their reprogramming of functions. Second, we report DON-induced increase in the intracellular glutamine metabolites occurring in the M1 macrophages; whether this DON-induced metabolic program is restricted to M1 macrophages and myeloid cells or if it also affects tumor cells and T cells remains to be determined. Additionally, while enhanced phagocytosis may be a potential mechanism driving the increased glutamine and its downstream metabolites in M1 macrophages, the exact mechanism and whether it is context-dependent entail further investigation. Finally, the current findings report that the differential regulation of M1 and M2-like macrophages by DON treatment can shed light on future glutamine metabolism-based strategies for cancer therapy.

## Declarations

### Availability of data and materials

The datasets used and/or analyzed during the current study are available from the corresponding author upon reasonable request.

### Author contributions

SQ, JF, and TLW designed the experiments. SQ, JF, and DSW conducted the experiments and performed data analyses. SQ, JF, DSW, SG, IMS, and TLW wrote the paper.

## Acknowledgments

The authors appreciate the initial help and guidance provided by Juliane Libert.

## Supplemental information

Figures S1–S7

Tables S1-S3

STAR Table

