## Supplemental Data 1 for "Modulating Glutamine Metabolism Reprograms Pro-Inflammatory Differentiation in Macrophages"

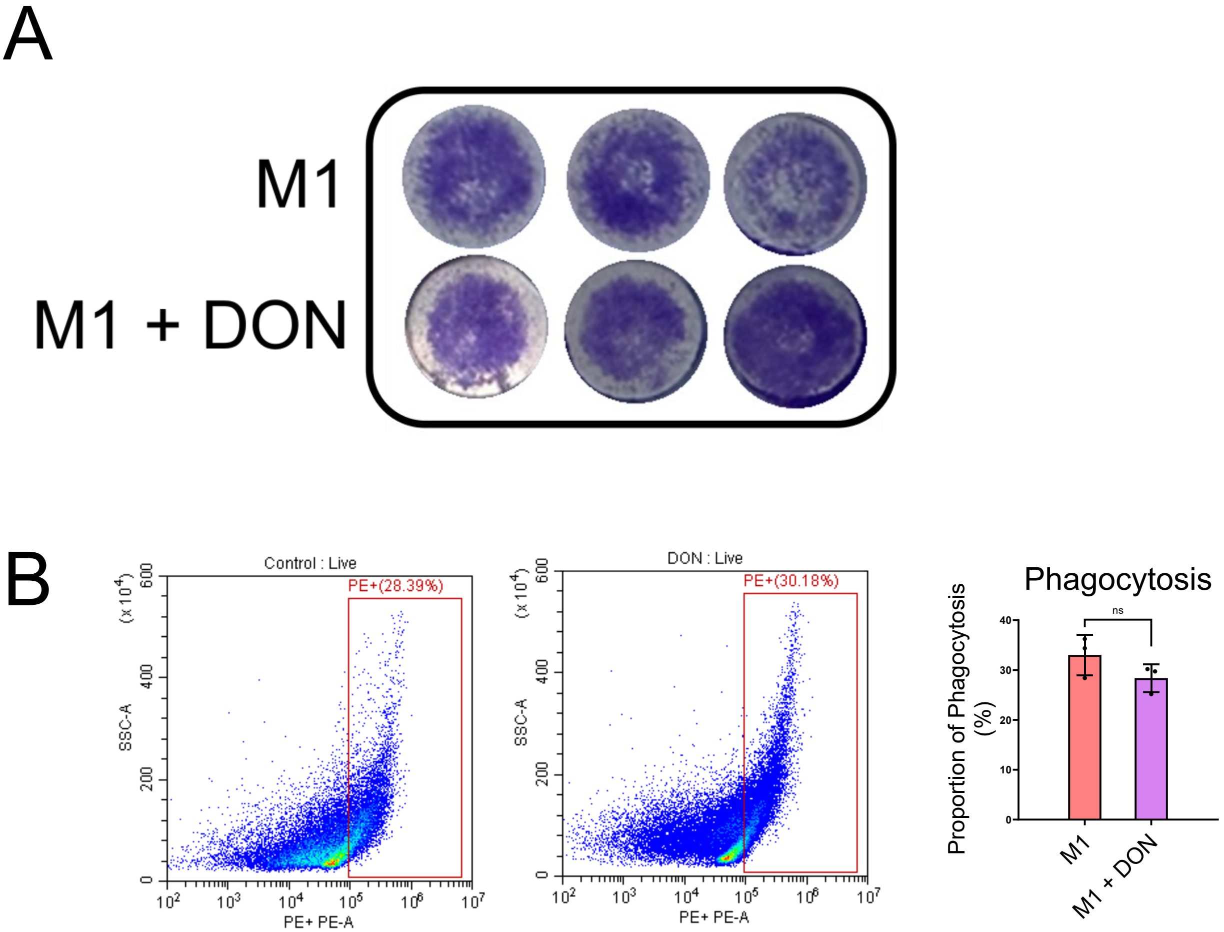


**Supplementary Figure 1. Pro-Inflammatory Functions of Full-Polarized M1 Macrophages**
**(A)** Transwell invasion assay of ID8 ovarian cancer cells co-cultured with M1-differentiated BMDMs that had been treated with or without DON using full-polarized protocols. There was no DON during the co-culture period.
**(B)** Figures illustrate representative flow cytometry plots (left) and phagocytic cell quantification (right).

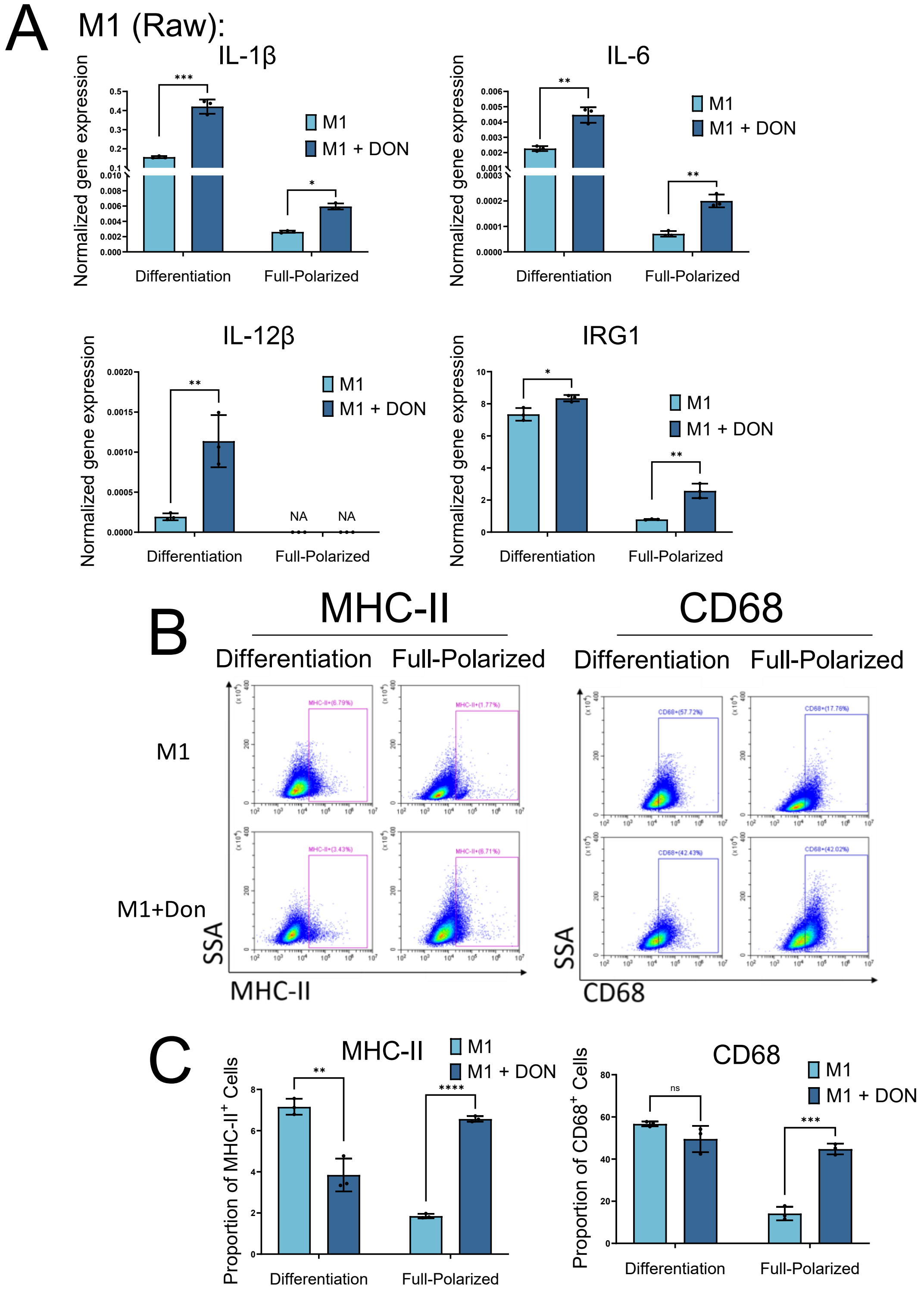


**Supplementary Figure 2. Effects of DON on M1-associated genes and surface markers in Raw 264.7 cells**

**(A)** mRNA expression of M1-associated genes (IL-1β, IL-6, IL-12β, and IRG1) in Raw 264.7 cells were analyzed using qPCR during differentiation (24 hours) and full-polarized conditions, with or without DON treatment.

**(B)** Flow cytometry analysis of M1 surface markers (MHC-II and CD68) in Raw 264.7 cells under differentiation and full-polarized conditions.

**(C)** Quantification of the flow cytometry results shown in (B), indicating the percentage of MHC-II$^{+}$ and CD68$^{+}$ cells in each group.

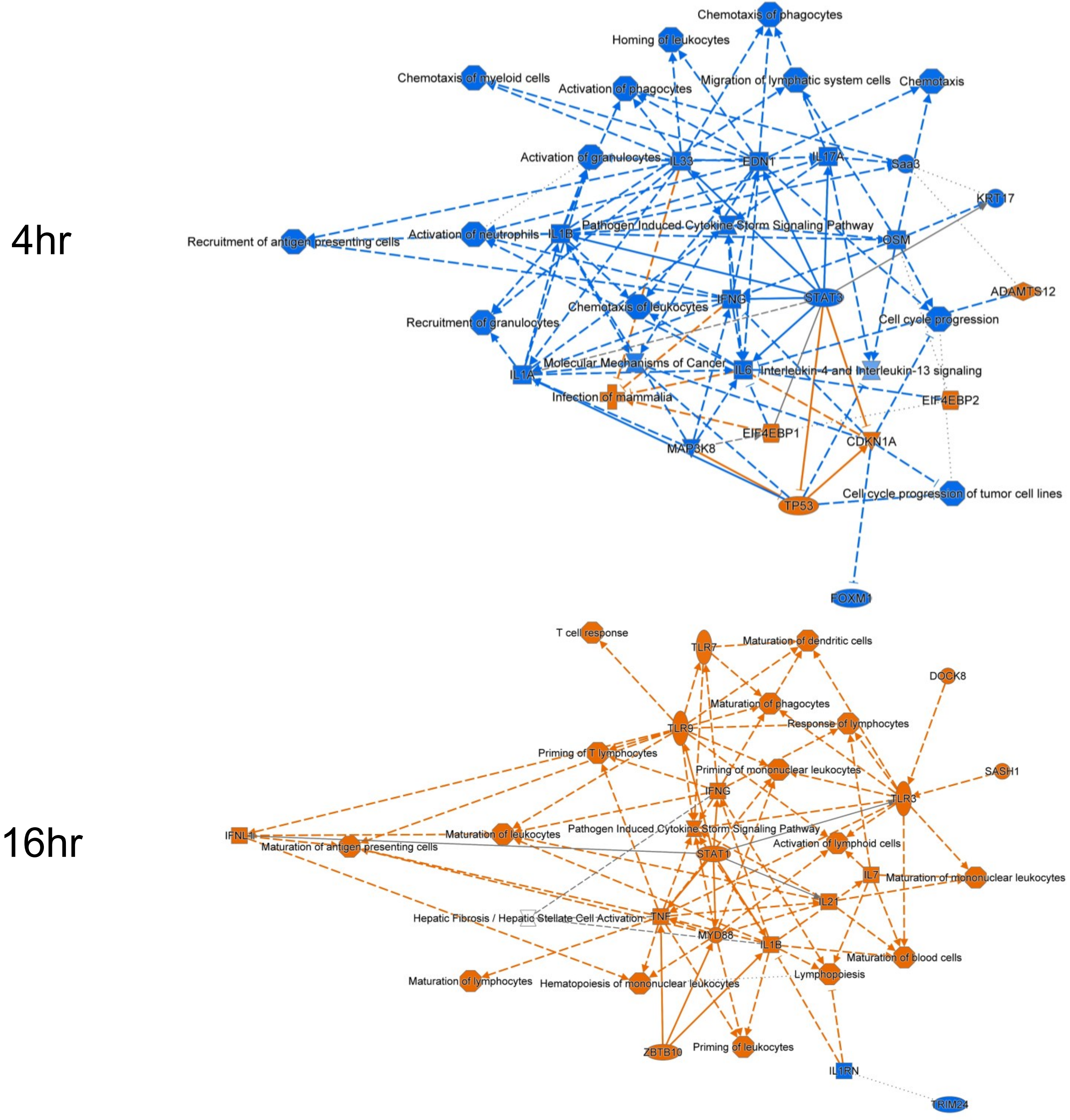


**Supplementary Figure 3. Upstream Regulator Predicted by the IPA Network Analysis**
Graphical summary of upstream regulator networks predicted by Ingenuity Pathway Analysis (IPA) for DON-treated versus control macrophages at 4 h (top) and 16 h (bottom). Predicted activation states are indicated by node color (orange: activated; blue: inhibited), while edge styles depict predicted regulator-target interactions.

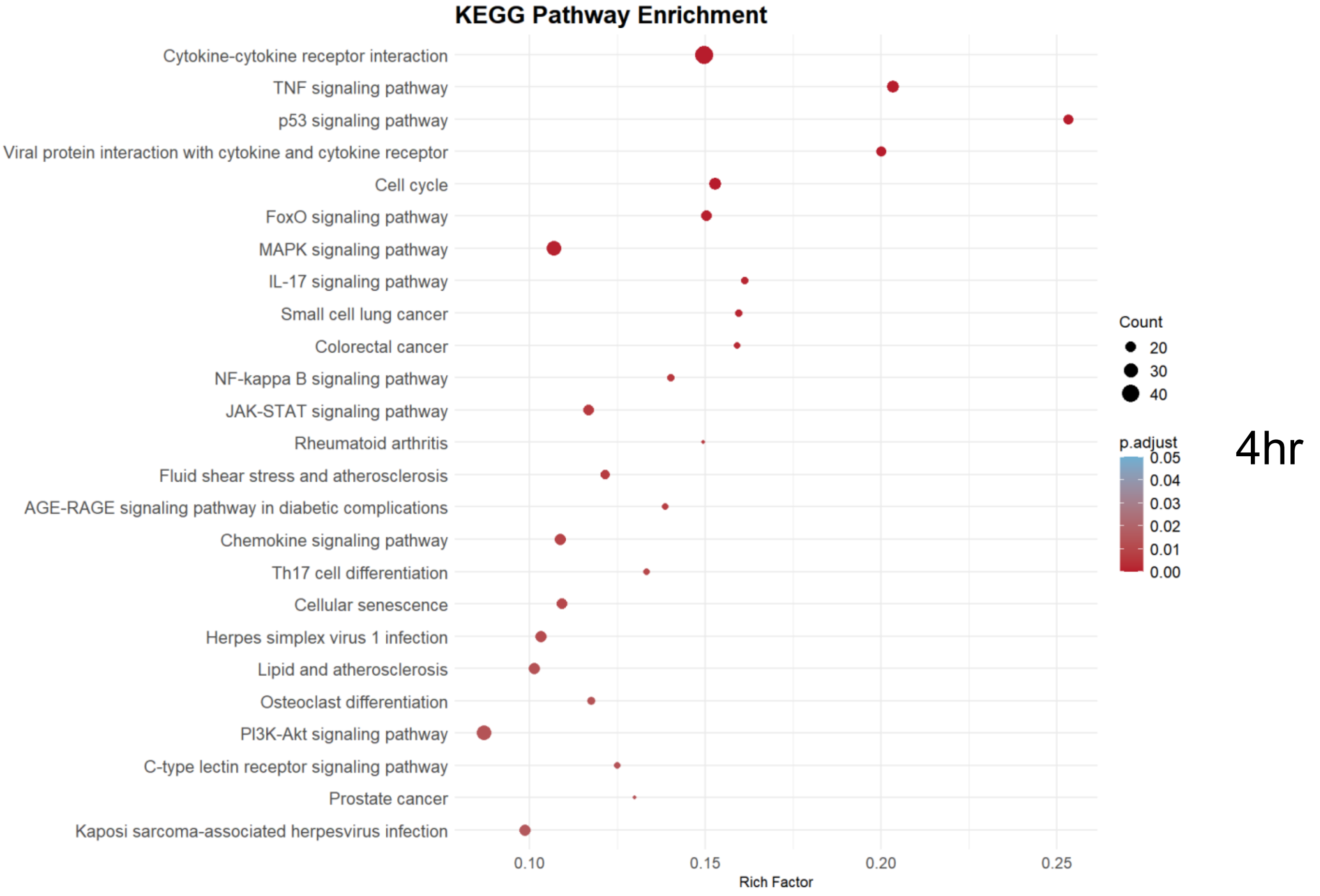
KEGG Pathway Enrichment
Cytokine-cytokine receptor interaction
TNF signaling pathway
p53 signaling pathway
Viral protein interaction with cytokine and cytokine receptor
Cell cycle
FoxO signaling pathway
MAPK signaling pathway
IL-17 signaling pathway
Small cell lung cancer
Colorectal cancer
NF-kappa B signaling pathway
JAK-STAT signaling pathway
Rheumatoid arthritis
Fluid shear stress and atherosclerosis
AGE-RAGE signaling pathway in diabetic complications
Chemokine signaling pathway
Th17 cell differentiation
Cellular senescence
Herpes simplex virus 1 infection
Lipid and atherosclerosis
Osteoclast differentiation
PI3K-Akt signaling pathway
C-type lectin receptor signaling pathway
Prostate cancer
Kaposi sarcoma-associated herpesvirus infection
0.10
0.15
0.20
0.25
Rich Factor
Count
20
30
40
p.adjust
0.05
0.04
0.03
0.02
0.01
0.00
4hr

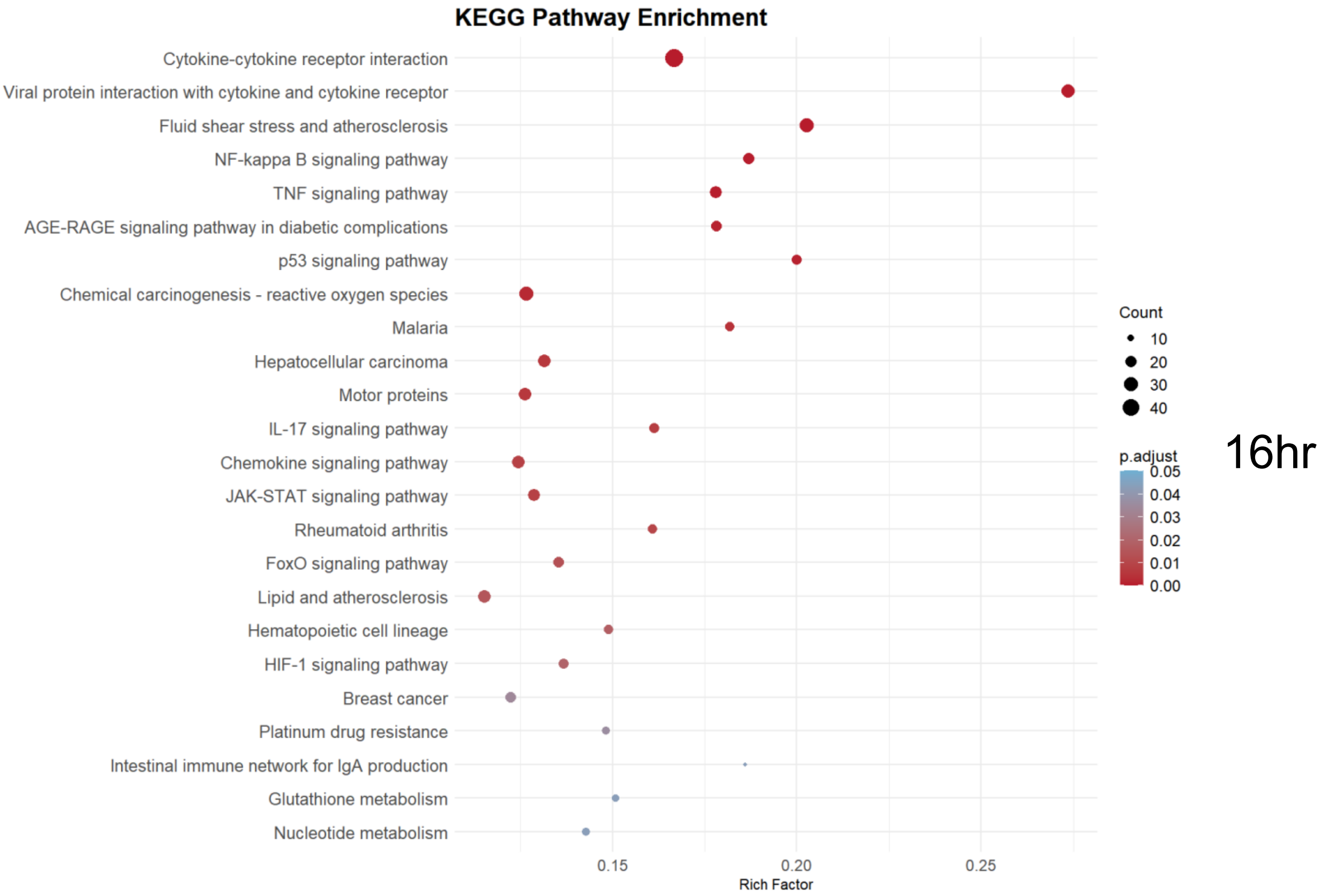
KEGG Pathway Enrichment
Cytokine-cytokine receptor interaction
Viral protein interaction with cytokine and cytokine receptor
Fluid shear stress and atherosclerosis
NF-kappa B signaling pathway
TNF signaling pathway
AGE-RAGE signaling pathway in diabetic complications
p53 signaling pathway
Chemical carcinogenesis - reactive oxygen species
Malaria
Hepatocellular carcinoma
Motor proteins
IL-17 signaling pathway
Chemokine signaling pathway
JAK-STAT signaling pathway
Rheumatoid arthritis
FoxO signaling pathway
Lipid and atherosclerosis
Hematopoietic cell lineage
HIF-1 signaling pathway
Breast cancer
Platinum drug resistance
Intestinal immune network for IgA production
Glutathione metabolism
Nucleotide metabolism
0.15
0.20
0.25
Rich Factor
Count
10
20
30
40
p.adjust
0.05
0.04
0.03
0.02
0.01
0.00
16hr

**Supplementary Figure 4. KEGG Pathway Analysis of Bulk RNA-seq Data**
KEGG pathway enrichment analyses using differentially expressed genes from bulk RNA-seq at 4 h (up) and 16 h (down) following DON treatment.

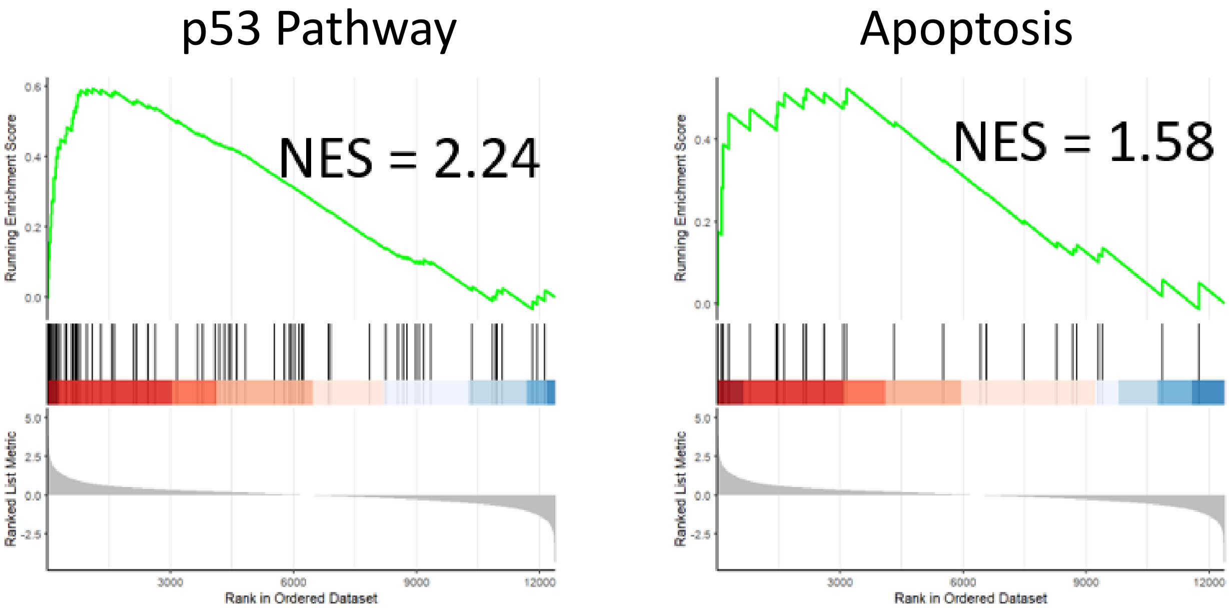


**Supplementary Figure 5. Gene Set Enrichment Analysis of p53 and Apoptosis Pathways in M1 Differentiated Cells.**
Gene Set Enrichment Analysis (GSEA) was performed on DEGs at 4 hours of DON or vehicle treatment. Normalized Enrichment Score (NES) and adjusted p-values (P.adjust) are as follows: p53 Pathway: NES = 2.24, P.adjust = 2.84e-07; Apoptosis: NES = 1.58, P.adjust = 0.026

Differentiation (4 Hours):

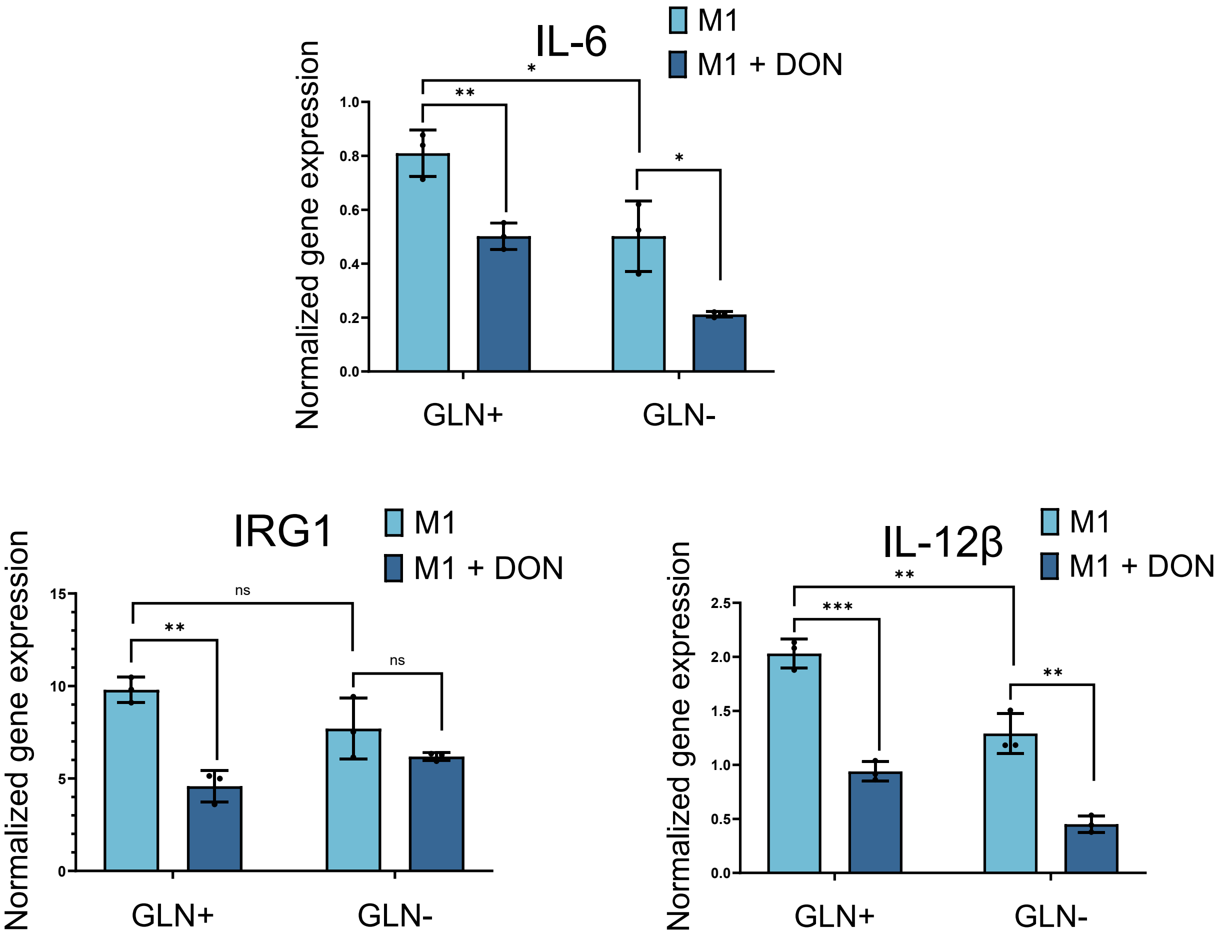


**Supplementary Figure 6. Effect of DON on M1-Associated Genes and Surface Markers in Glutamine-Replete ($GLN^+$) and Glutamine-Depleted ($GLN^-$) Conditions in the 4-Hour Differentiation Model.** qPCR analysis of M1-associated gene expression (IL-6, IL-1β, and IRG1) in glutamine-replete ($GLN^+$) or glutamine-depleted ($GLN^-$) conditions. Study was performed on M1-differentiation model (4 hr).

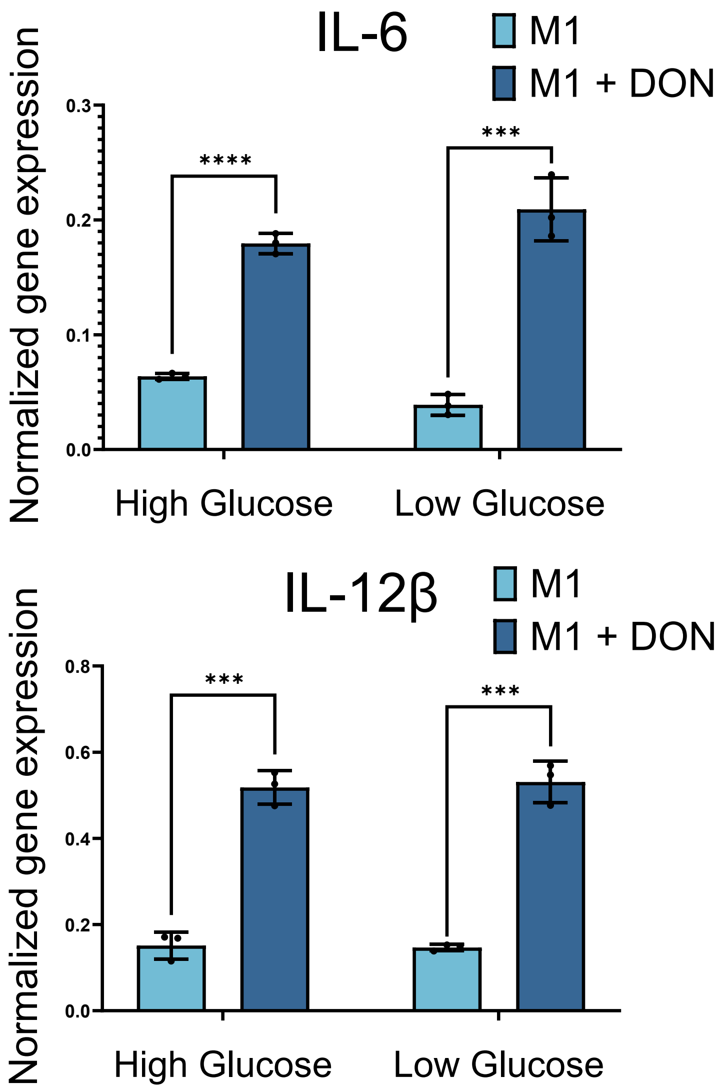


**Supplementary Figure 7. The effect of Extracellular Glucose Availability on DON-Induced M1 Macrophage Activation.**
qPCR analysis of IL-6 and IL-12 gene expression in M1 macrophages differentiated for 16 hours under high-glucose (HG) or low-glucose (LG) conditions, with or without DON (10 μM) treatment.
